# Changes in maternal blood and placental lipidomic profile in obesity and gestational diabetes: Evidence for sexual dimorphism

**DOI:** 10.1101/2024.07.24.605016

**Authors:** Leena Kadam, Marija Veličković, Kelly Stratton, Carrie D. Nicora, Jennifer E. Kyle, Eric Wang, Matthew E. Monroe, Lisa M. Bramer, Leslie Myatt, Kristin E. Burnum-Johnson

**Author notes:** Corresponding author, 3303 S Bond Avenue, CHH1, OHSU, Portland, OR 97239.

## Abstract

**Introduction:** Obesity and gestational diabetes (GDM) are associated with adverse pregnancy outcomes and program the offspring for cardiometabolic disease in a sexually dimorphic manner. The placenta transfers lipids to the fetus and uses these substrates to support its own metabolism impacting the amount of substrate available to the growing fetus.

**Methods:** We collected maternal plasma and placental villous tissue following elective cesarean section at term from women who were lean (pre-pregnancy BMI 18.5-24.9), obese (BMI>30) and type A2 GDM (matched to obese BMI) with male or female fetus (n=4 each group). Lipids were extracted and fatty acid composition of different lipid classes were analyzed by LC-MS/MS analysis. Significant changes in GDM vs obese, GDM vs lean, and obese vs lean were determined using t-test with a Tukey correction set at p<0.05.

**Results:** In placental samples 436 lipids were identified, among which 85 showed significant changes. Of note only in male placentas significant decreases in C22:6 - docosahexaenoic acid (DHA) in phosphatidylcholine (PC) and triglyceride lipid species were seen when comparing tissue from GDM women to lean. In maternal plasma we observed no effect of obesity. GDM or fetal sex.

**Conclusion:** This is the first study assessing fatty acid composition of lipids in matched maternal plasma and placental tissue from lean, obese, and GDM women stratified by fetal sex. It highlights how GDM affects distribution of fatty acids in lipid classes changes in a sexually dimorphic manner in the placenta.

## INTRODUCTION

The obesity epidemic (BMI>30) is an escalating threat to population health and healthcare systems worldwide. More than 1/3^rd^ women in the United States are obese and one half of pregnant women are overweight or obese [1]. Compared to age-matched lean women (pre-pregnancy BMI 18.5-24.9), obese women have increased risk of spontaneous abortion, recurrent miscarriage, and stillbirth particularly with a male fetus [2-4]. Obesity also increases the risk of almost all complications of pregnancy including gestational diabetes mellitus (GDM) [5-8] as obese women are 4 times more likely to develop GDM [9, 10]. GDM manifests as impaired glucose tolerance from the late 2^nd^ trimester of pregnancy and has an increasing prevalence, ∼7% of pregnancies, in the US [11]. Women with obesity and GDM display hyperglycemia and hyperlipidemia; are more susceptible to hypertensive disorders; dysfunctional labor; thromboembolic events and are at increased risk of developing type 2 diabetes in later life [1, 12-15]. Fetal and newborn complications in these pregnancies include congenital malformations, large-for-gestational-age (LGA) infants, intrauterine growth restriction (IUGR), cesarean delivery, birth trauma, asphyxia, and stillbirth; and importantly, programming for subsequent development of obesity and diabetes [1, 9, 12, 16-20]. The fetal programming effect of obesity and GDM contributes to the vicious cycle of their development, with women carrying a male fetus being reportedly at a higher risk to develop adverse pregnancy outcomes underscoring the sexual dimorphic effects of these conditions.

Differences in lipid profiles and metabolism have been previously described for obesity and diabetes in non-pregnant individuals [21, 22]. Several studies have explored lipid composition of maternal and corresponding cord plasma (fetal) in obesity and GDM with contrasting results [23-28]. Significant increase in maternal fatty acids (FA) in the 3^rd^ trimester in GDM vs controls with alterations in specific FA [23-26] [28] have been reported. Studies of cord blood fatty acids, particularly DHA, reveal differences in lipid FA composition correlating with fatty acid transporter activity or handling/metabolism of fatty acids by the fetus [23] [29]. Interestingly, none of these early studies considered the effect of fetal sex or placental fatty acid metabolism. During pregnancy, the placenta regulates maternal metabolism to ensure supply of nutrients to support both its own metabolism and transfer to the fetus. At term the placenta is 1/6^th^ the size of the fetus but, has 6 times the metabolic rate and changes in placental lipid metabolism may influence the amount that reaches the fetus, potentially altering fetal growth. Placental exposure to excess maternal lipids seen in obesity and GDM, is reported to alter metabolic pathways involved in energy storage, FA oxidation, inflammation, cell growth, death, and differentiation in animals and humans [30-34]. FA’s are important nutrients necessary for normal growth and development of the fetus and their supply is particularly critical in the 3^rd^ trimester of pregnancy due to rapid fetal growth and nutrient demand [35-37]. Furthermore, as components of triglycerides, phospholipids and other complex lipids, FA’s perform several crucial cellular functions including maintaining membrane integrity, signal transduction, cellular metabolism, energy storage, etc. The hyperlipidemia and hyperglycemia seen in obesity and GDM can thus affect the uptake and metabolism of FA’s by the placenta and what gets transferred to the fetus [38]. Understanding the changes in placental lipid metabolism and distribution of FA’s can offer mechanistic insights into the effects of GDM and obesity on maternal-fetal health.

There is abundant evidence that fetal sex affects *in-utero* growth strategy with the male having a more aggressive strategy than the female, but also birth and neonatal outcomes as well as programming of adult disease [39-42]. We have previously demonstrated sexual dimorphism in trophoblast mitochondrial respiration, in placental autophagy, and in placental expression of antioxidant enzymes with obesity and GDM [43-45]. However, very few studies have explored sexual dimorphism in placental lipid metabolism and how it changes across the maternal-placental-fetal compartments during obesity and GDM. Understanding how maternal factors, via the placenta, affect the developing fetus in a sex-specific manner is important to the development of prevention strategies for chronic disease.

In this study we used a lipidomic approach to test the hypothesis that we would find asexual dimorphism in the FA composition of different lipid classes in matched maternal plasma and placental tissue from lean, obese, and BMI-matched type A2 GDM women.

## MATERIALS AND METHODS

### Sample collection

Informed consent was obtained from all patients under a protocol approved by the Institutional Review Board of Oregon Health & Science University, prior to the tissue being de-identified and placed in a tissue repository. Maternal blood and placental tissue were collected from three groups of women: Lean (pre-pregnancy BMI 18.5-25.0), Obese (pre-pregnancy BMI 30.1-45.0) and type A2GDM (GDM) (BMI matched to obese group) with either male or female fetus (n=8 each group and n=4 per sex). GDM was defined using IADPSG criteria which recommends 75-grams two hours OGTT with at least one abnormal result: Fasting plasma glucose (FPG) at ≥ 92 mg/dl or 1-hour ≥ 180 mg/dl or 2-hour ≥ 153mg/dl to be classified as GDM [46]. Women with type A2 GDM were those with greater than 30% of FPG ≥90-95 mg/dl or 1 hr Postprandial ≥130-140 mg/dl and needed medication to control their blood glucose levels. To keep the samples consistent, we only collected samples from women who received insulin as medication. Inclusion criteria included a singleton pregnancy, an age range of 18-45 and delivery by cesarean with no labor. Exclusion criteria included concurrent disease (including, but not limited to IUGR, hypertension, pre-eclampsia, eating disorders, infection, inflammatory disorders), use of tobacco, drugs, or medications other than to treat GDM, excessive weight gain or loss prior to pregnancy (>20 lbs), or bariatric surgery in the last year and labor defined by regular uterine contractions (every 3-4 minutes, verified by tocodynamometry) resulting in cervical dilatation and/or effacement.

Maternal blood was collected from fasting patients before elective C-section and placed in EDTA-containing collection tubes. Plasma was separated by centrifugation at 2000 g for 10 min at 4°C, then flash-frozen in liquid nitrogen and stored at −80°C. Placentas were collected post-delivery and villous tissue (VT) was immediately sampled from 5 random sites of the placenta, flash frozen and stored at -80^0^C. Subsequently the villous tissue was powdered under liquid nitrogen and equal amounts of the 5 separate samples combined before analysis for each subject.

### Lipid extraction

One mL of an ice-cold 2:1 (v/v) chloroform:methanol mixture was pipetted into a 2 mL Sorenson MulTI™ SafeSeal™ microcentrifuge tubes (Sorenson bioscience, Salt Lake City, UT) inside an ice-block. 200 μl of the maternal plasma was then added to each of the Sorenson tubes at a ratio of 1:5 sample:chloroform/methanol mixture and vigorously vortexed. The liquid nitrogen ground and powdered placental tissue was also added to 2 mL Sorenson tubes along with 200 μl of water and 1 mL of the 2:1 (v/v) chloroform:methanol solution. The placental tissue was homogenized with a handheld pestle motor mixer with spiraled pestles (Cole-Palmer, Vernon Hills, IL). The samples were placed in the ice block for 5 mins and then vortexed for 10 secs followed by centrifugation at 10,000 xg for 10 mins at 4°C. The lower lipid soluble phase was transferred to a glass vial for lipidomics analysis, and dried in a speed vac. 500 μl of 2:1 (v/v) chloroform:methanol was added to the glass vial, capped and stored at -20°C.

### Lipid analysis

Samples were analyzed using liquid chromatography-tandem mass spectrometry (LC-MS/MS). Briefly, the dried lipid extracts were reconstituted in methanol and10 μl was injected onto a reversed-phase Waters CSH column (3.0 mm × 150 mm × 1.7 μm particle size) connected to a Waters Aquity UPLC H class system interfaced with a Velos-ETD Orbitrap mass spectrometer. Lipid molecular species were separated over a 34 min gradient using mobile phase A (acetonitrile/water (40:60) containing 10 mM ammonium acetate) and mobile phase B (acetonitrile/ isopropyl alcohol (10:90) containing 10 mM ammonium acetate) at a flow rate of 250 μl/min.

Samples were analyzed in both positive and negative ionization using HCD (higher-energy collision dissociation) and CID (collision-induced dissociation) to obtain high coverage of the lipidome. Confident lipid identifications were made using in-house developed identification software LIQUID [47]. To facilitate the quantification of lipids, a reference database for lipids identified from the MS/MS data was created and features from each analysis were then aligned to the reference database based on their identification, m/z, and retention time using MZmine 2 [48]. Aligned features were manually verified and peak apex intensity values were exported for subsequent statistical analysis.

### Statistical Methods

The data from both sample types was analyzed using the pmartR package[48, 49]. First, data were log2 transformed separately for negative and positive ionization mode, and potential outliers were identified via RMD-PAV with correlation, median absolute deviation, and skewness as the metrics utilized [50]. The decision of whether to remove a sample as an outlier was based on a holistic view of the RMD-PAV results in combination with Pearson correlation heatmap and probabilistic principal components analysis (PCA) plot. Data were normalized via global median centering, after which point negative and positive ionization modes were combined into a single dataset. A t-test with Tukey p-value adjustment was performed to compare each pair of maternal groups (GDM, obese, and lean), regardless of fetal sex. An additional t-test was carried out separately for the male fetus and female fetus placental samples (Supplementary Table 1, 2).

**TABLE 1.**
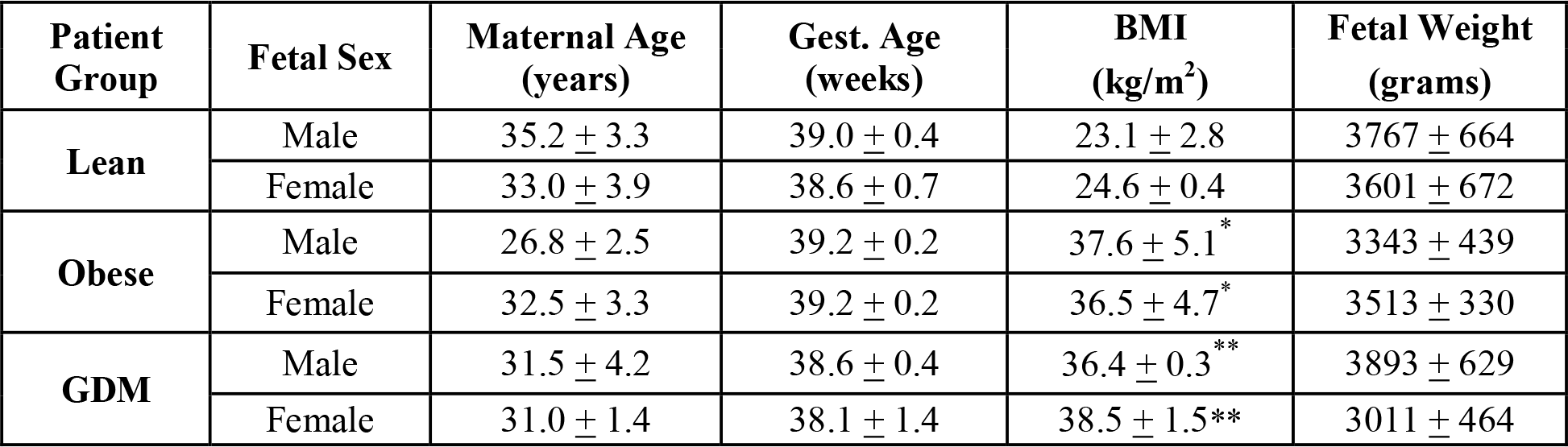
Patient Characteristics. Values are expressed as Mean ± SD, n=4 each group. * p< 0.005 vs. Lean, ** p<0.0001 vs. Lean of corresponding sex

## RESULTS

### Patient characteristics

Patients were matched for maternal age, gestational age at delivery, and the obese and GDM groups were matched for pre-pregnancy BMI (Table 1). This enabled us to identify the distinct lipid profile potentially resulting from hyperinsulinemia and hyperlipidemia in obesity and GDM. All neonates were born appropriate for gestational age (AGA) and there was no difference in fetal weight between lean, obese and GDM groups. Placental weights were not routinely recorded.

### Lipidomic analyses

In total 436 and 284 lipids were quantified in the placenta and maternal blood samples respectively, providing insight into the molecular dialog between the placental metabolism and maternal metabolic environment. We also assessed the differences in fatty acid composition in detected lipids. This discovery-based LC-MS/MS lipidomic approach provides a picture of how individual lipid species change in various tissues and across conditions and data is presented as heatmaps for each category of lipid classes with differences in their FA composition in each figure with individual plasma or placental samples from each patient separated into columns.

#### Placenta

In the placenta, 176 lipids were identified in negative ionization mode and 260 in positive. No potential outliers were identified in either ionization mode. In the combined male and female analysis, there were 85 statistically significant lipids (Tukey adjusted p-value < 0.05 for at least one of the three comparisons) identified across both ionization modes. When sex stratified for female and males placentas, samples there were 17 and 73 significant lipids (Tukey adjusted p-value < 0.05 for at least one of the three comparisons) respectively. Placental sample data and statistical results are listed in Supplementary Table 1.

We observed significant differences in the triacylglycerols (TG), diacylglycerides (DG), phosphatidylcholines (PC), phosphatidylinositol (PI), phosphatidylglycerols (PG), phosphatidylethanolamines (PE), phosphatidylserine (PS), Ceramides (Cer) and sphingomyelins (SM) many of which contained at least one long-chain polyunsaturated fatty acid (LCPUFA). Data are discussed for each class of lipids.

##### Triacylglycerol’s (TG)

We observed a significantly lowered levels of TG’s containing combinations of C16:0 (palmitic acid), C18:0 (stearic acid), C22:5 (docosapentaenoic acid), C22:6 (docosahexaenoic acid, DHA) and C20:4 (arachidonic acid, AA) across their three FA positions in GDM placentae compared to Lean when both sexes were analyzed together (Figure 1A). DHA and AA are essential FA’s that cannot be synthesized in the placenta and obtained from the mother. All the 16 TG species downregulated in GDM (v/s Lean) placentae had at least one DHA in them and 7 had AA. Two TG’s were elevated in Obese placentae (vs Lean) both with DHA in at least one position (Figure 1A). When stratified by fetal sex, we observed this decrease was mainly driven by placentae from male fetuses, with significantly decreased levels in 13 of the total 16 species observed in combined analysis and 1 unique TG (TG 20:4_22:5_22:6) also containing DHA). Of the 13, 6 TG’s were also significantly lower in GDM vs obese comparison in addition to TG (20:4_22:5_22:6) and TG (18:0_20:0_22:6) which were not observed in the combined analysis. We also observed increased levels of TG species containing 51-64 total carbons in GDM placentas (vs Lean) but were not able to confidently resolve their constituent FA’s (Supplementary Figure).

**Figure 1.**
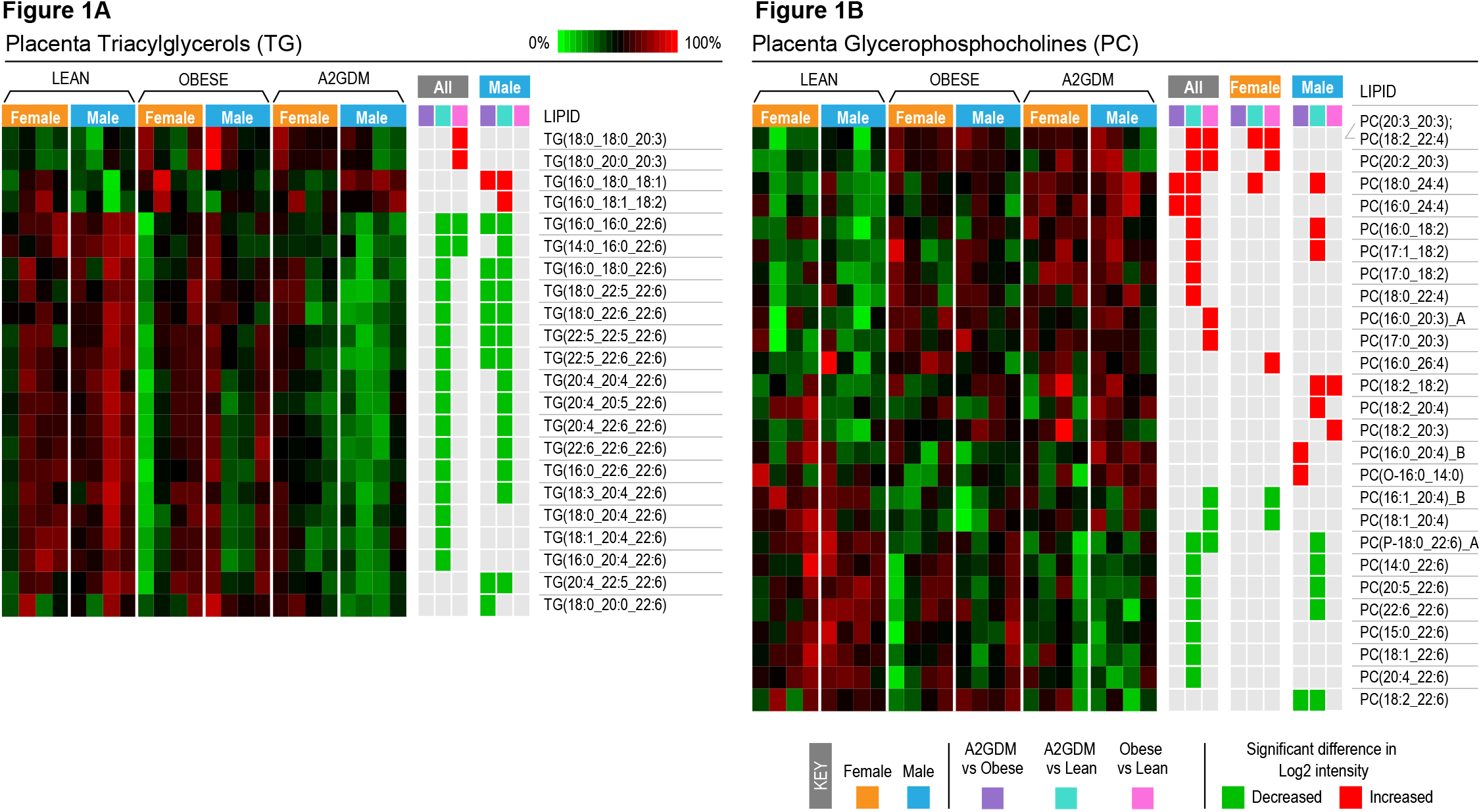
Heatmaps representing the relative log2 expression levels of lipids in **A:** Triacylglycerols (TG) and **B:** Glycerophosphocholines (PC) in placental tissue of lean, obese and GDM women with either a male or a female fetus (n=4 each group). Significant increases or decreases in specific lipid composition (adjusted p<0.05, t-test with Tukey’s comparison) between GDM vs obese, GDM vs lean and obese vs lean groups for all women combined or those with a male or a female fetus. Each column represents an individual patient sample, and each row represents the relative abundance of a unique lipid, and the heatmaps are scaled by row. Lipid abbreviations for each acyl chain (number of carbons:number of double bonds). Structural isomers are denoted with _A or _B.

##### Phosphatidylcholine (PC)

We observed significant changes in PC levels in the bit sex combined and stratified. In GDM vs Lean comparison, we observed significant increase in PC containing combinations of palmitic acid, stearic acid, DGLA, linoleic acid and its derivatives and very long chain FA’s (C24) linked to the glycerol backbone. We observed significantly decreased levels of PC predominantly containing DHA which were maintained in male GDM (vs lean males) (Figure 1B). In the combined analysis of GDM vs Obese, two PCs both containing very long chain FA (PC18:0_24:4 and PC16:0_24:4) were increased. In the Obese vs Lean comparison, PC containing derivatives of dihomo-γ-linolenic acid (C20:3 or DGLA) were significantly increased and those containing DHA and AA were significantly decreased. Sex stratified analysis showed that these trends were driven by females (Figure 1B).

##### Phosphinositol (PI)

We observed significantly decreased levels of palmitic acid, stearic acid, oleic acid, DGLA, DHA and AA in PI in GDM placenta (vs Lean). Stratifying the data based on fetal sex showed that only males had significantly reduced levels of PI with DHA (Figure 2A).

**Figure 2.**
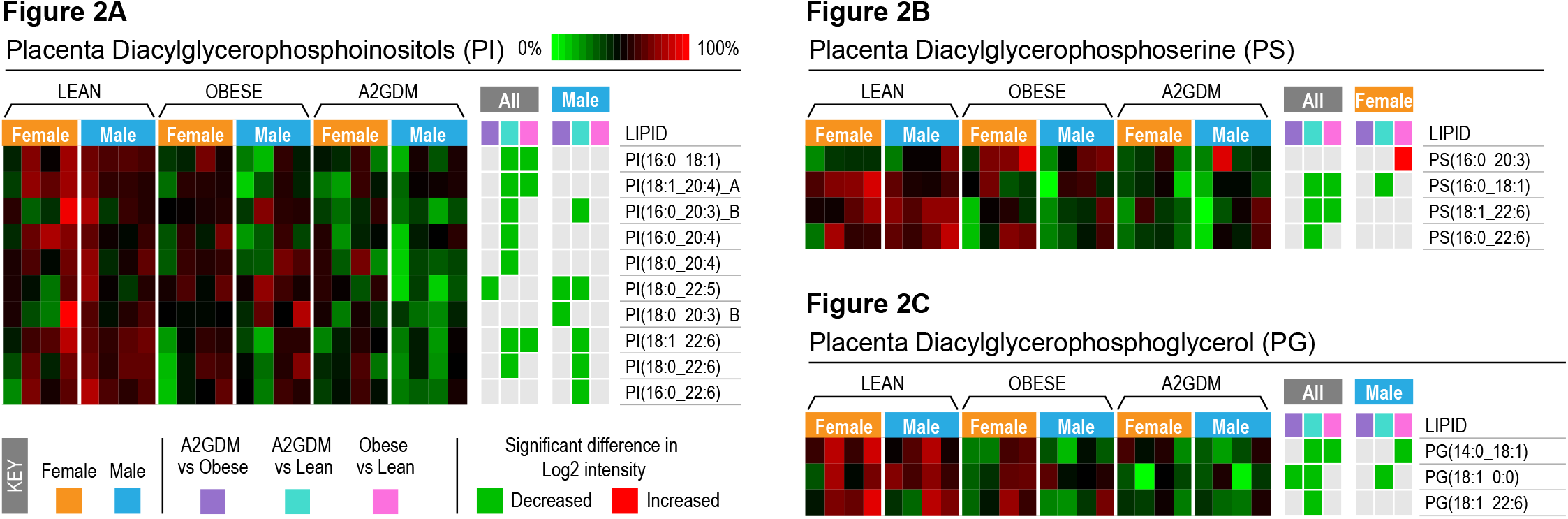
Heatmaps representing the relative log2 expression levels of **A:** Diacylglycerophosphoinositol (PI), **B:** Diacylglycerophosphoserine (PS), and **C:** Diacylglycerophosphoglycerol (PG) lipids in placental tissue of lean, obese and GDM women with either a male or a female fetus (n=4 each group). Significant increases or decreases in specific lipid composition (adjusted p<0.05, t-test with Tukey’s comparison) between GDM vs obese, GDM vs lean and obese vs lean groups for all women combined or those with a male or a female fetus. Each row represents the relative abundance of a unique lipid, and the heatmaps are scaled by row. Lipid abbreviations show the total number of acyl chain carbons:total number of double bonds. Structural isomers are denoted with _A or _B.

##### Phosphatidylserines (PS)

In the sex combined analysis, GDM placentae had significantly reduced levels of 3 out of 4 differential lipids detected and contained palmitic acid, oleic acid and DHA when compared to Lean and none with Obese. In sex stratified analysis, only females showed significant differences with female GDM having lower levels of PS (16:0_18:1) and obese samples showing increased level of PS (16:0_20:3) (Figure 2B) both compared to Lean.

##### Phosphatidylglycerol (PG)

Decreased levels of PG (14:0_18:1), (18:1_22:6) and (18:1_0:0) were observed in GDM placentae from both male and female fetuses when compared to Lean. Sex stratified analysis showed that these were driven by males (Figure 2C).

##### Phosphoethanolamines (PE)

Significantly elevated levels of 15 PE species containing combinations of palmitic acid, stearic acid, linoleic acid, DGLA and AA but increased levels of PE with DHA (22:6_22:6)were found in GDM placentae (vs Lean) (Figure 2A). Sex stratified analysis showed that this was driven by males in addition to 6 more PE. Both obese & GDM females had significantly reduced levels of DHA vs Leans. (Figure 3A).

**Figure 3.**
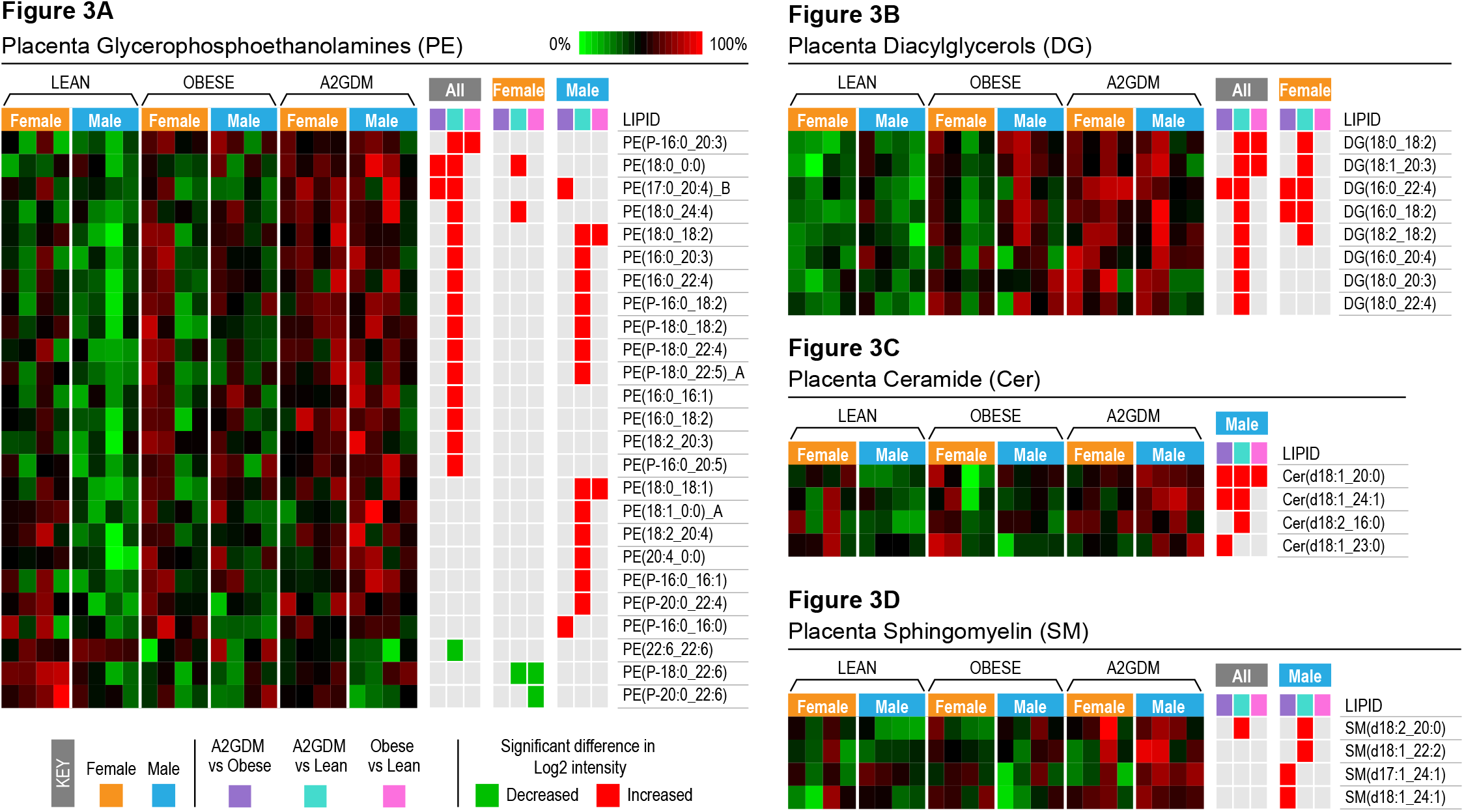
Heatmaps representing the relative log2 expression levels of **A:** Glycerophosphoethanolamine (PE), **B:** Diacylglycerol (DG), **C:** Ceramide (Cer), and **D:** Sphingomyelin (SM) lipids in placental tissue of lean, obese and GDM women with either a male or a female fetus (n=4 each group). Significant increases or decreases in specific lipid composition (adjusted p<0.05, t-test with Tukey’s comparison) between GDM vs obese, GDM vs lean and obese vs lean groups for all women combined or those with a male or a female fetus. Each row represents the relative abundance of a unique lipid, and the heatmaps are scaled by row. Lipid abbreviations show the total number of acyl chain carbons:number of double bonds. Structural isomers are denoted with _A or _B.

##### Diacylglycerols (DG)

In the sex combined analysis, we observed significant increase in 8 DG containing combinations of palmitic acid, stearic acid, linoleic acid, DGLA and C22:4 (adrenic acid, a source for AA) in GDM (vs Lean) (Figure 3B). Of these 8, 2 were also high in Obese (vs Lean) and 1 in GDM (vs Obese). Sex stratified analysis showed that these differences were driven by females, with 5 of 8 detected DG elevated in GDM placentae (vs Lean) and 2 in Obese (vs GDM). female

##### Ceramides

No significant differences were observed in the sex combined analysis. When stratified by sex, increased levels of ceramides were observed only in male across all three comparisons (Figure 3C).

##### Sphingomyelin (SM)

Significant differences were observed in 2 SM in the male GDM placentae (vs Lean) with only 1 elevated in the sex combined analysis. GDM placentae had elevated levels of wo SM both with C24:1 (nervonic acid) chain elevated when compared to Obese group (Figure 3D).

#### Maternal Plasma

In the maternal plasma samples, 64 lipids were identified in negative ionization mode and 220 in positive. One potential sample outlier was identified via RMD-PAV in negative ionization mode but retained in the dataset after considering the correlation and PCA results. In the sex combined analysis, there were 14 statistically significant lipids (Tukey adjusted p-value < 0.05 in at least one of the three comparisons). Maternal sample data and statistical results are listed in Supplementary Table 2. We did not observe any effect of fetal sex in the maternal plasma data, and therefore report only the sex combined data analysis (Fig 4A-D).

**Figure 4.**
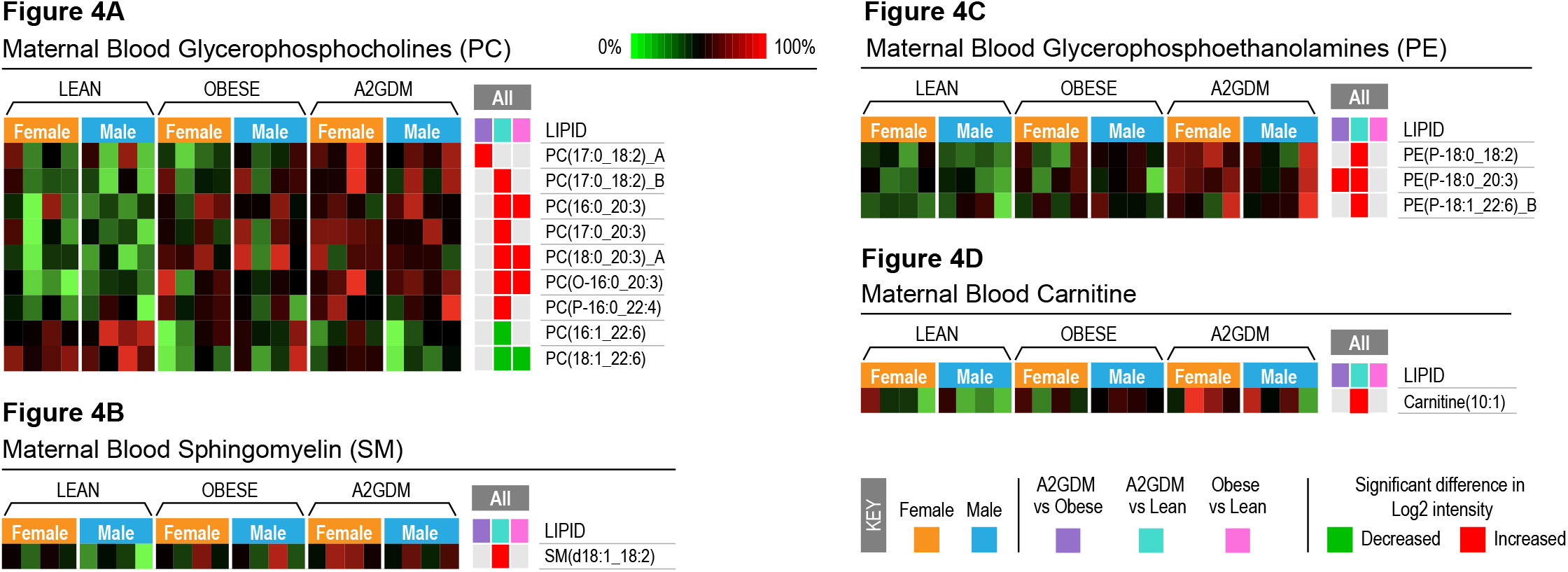
Heatmaps representing the relative log2 expression levels of **A:** Glycerophosphocholines (PC), **B:** Sphingomyelin (SM), **C:** Glycerophosphoethanolamine (PE), and **D:** Carnitine lipids in maternal blood of lean, obese and GDM women for male and female fetuses combined. Significant increases or decreases in specific lipid composition (adjusted p<0.05, t-test with Tukey’s comparison) between GDM vs obese, GDM vs lean and obese vs lean groups for all women combined. Each row represents the relative abundance of a unique lipid, and the heatmaps are scaled by row. Lipid abbreviations show the total number of acyl chain carbons:number of double bonds. Structural isomers are denoted with _A or _B

##### Phosphatidylcholine (PC)

Are the largest number of differentially expressed lipids observed in maternal plasma. GDM group showed increased levels of 6 PC species compared to lean, 4 of which contained DGLA and 1 had linoleic acid - both considered essential fatty acids for humans and decreased levels of two PC both containing DHA. Compared to obese, GDM had elevated levels of PC (17:0_18:2). The obese group had higher levels of 3 PC’s all containing DGLA when compared to Lean and and lower levels of 1 PC containing DHA, which overlapped with the GDM group (Figure 4A).

##### Sphingomyelin (SM)

Significantly increased levels SM containing linoleic acid SM(d18:1/18:2) was observed in GDM plasma when compared to Lean. No differences were seen in the GDM vs Obese or Obese vs Lean comparisons (Figure 4B).

##### Phosphoethanolamines (PE)

Significantly increased levels of 3 PE containing combination of stearic acid with linoleic acid or, DGLA, and oleic acid with DHA were observed in GDM vs Lean and the one stearic acid and DGLA was elevated in the GDM vs obese comparison (Figure 4C).

##### Carnitine

Were uniquely identified as significant in the maternal plasma with Carnitine (10:1) showing *elevated* levels in the GDM vs Lean comparison GDM samples with (Figure 4D).

## DISCUSSION

Here we aimed to understand distribution of FA’s incorporated into different lipids changes in placenta and maternal plasma in obesity and GDM. We applied an untargeted LC-MS/MS approach to determine the FA composition of 9 lipid classes in samples from lean, obese and GDM women, and examined how they are influenced by fetal (placental) sex. Compared to maternal plasma, placenta showed more differences which were sexually dimorphic. We identified 436 individual lipids in placental tissue with 85 lipids spread across 9 lipid classes being significantly different among the three clinical conditions.

Several studies have investigated lipid composition of maternal plasma in obesity and GDM often with contrasting results. Sobki *et al* reported no differences in maternal blood lipids (low-density lipoprotein (LDL), high density lipoprotein (HDL), total cholesterol (TC), TG, and total lipids) between Lean and GDM patients [19]. Ryckman *et al* in their meta-analysis of 60 studies reported elevated levels of TG in maternal blood in GDM but no differences in HDL, TC or LDL [27]. These studies focused on traditional lipid indicators (HDL and LDL) and utilized a targeted approach and did not assess FA composition. Studies utilizing an untargeted approach have focused on samples collected early in pregnancy before the diagnosis of GDM to determine an association or detect biomarker signatures (see [51] for a comprehensive review) and thus offer no insight into FA metabolism or distribution. Chen *et al* reported increase in FA’s in fasting blood samples from GDM vs control women with a graded relationship of severity of hyperglycemia and concentration of individual 16:0, 18:1, 18:2, 18:3 and 22:6 fatty acids, in contrast to our observations [28]. Differences in time of sample collection (3^rd^ trimester vs early 2^nd^ trimester) and non-distinction of GDM groups (GDM controlled by only insulin vs both dietary/insulin/other medications) could account for these differences. Furse *et al* [24] in their untargeted approach found increases C16:0, C16:1, C18:0, C18:1 FA’s in diglycerides and TG in GDM and attributed this to de novo lipogenesis. Using a similar approach, Ortega-Senovilla *et al* reported reduced levels of 16:0, 18:1, 18:2, 18:3 in GDM and lower levels in cord blood [52]. Both these studies had BMI matched obese euglycemic women as their controls making it difficult to delineate the combined effects of obesity and GDM on maternal lipid distribution. Studies assessing matched maternal plasma and placenta samples and accounting for fetal (placental) sex are rare. Sexual dimorphism in placental gene expression and function is linked to differences in pregnancy outcomes, with a male fetus being at greater risk of an adverse outcome [53] [43] [40, 41] and assessing the effects of obesity and GDM from a fetal sex perspective is therefore important.

We observed that in the placenta PE, PC and TG had the highest differences between GDM and Lean groups, with few differences in Obese vs Lean and GDM vs Obese comparisons. PE and PC were elevated in GDM whereas TG decreased with male placentae driving these differences. PE and PC are the two most abundant membrane lipids, with PC accounting for 40–50% of total cellular phospholipids and PE (with cardiolipin) accounting for 30-45% of mitochondrial inner membrane lipids [54-56]. Both PC and PE are important for energy metabolism with abnormally high, or low cellular levels influencing energy metabolism in various organelles and reduction in PE is associated with increased mitochondrial membrane potential, and reduced complex I, IV activity and ATP production [54-56]. Their altered levels could contribute to the altered placental metabolism and mitochondrial function reported in GDM [57]. TG act as energy reserves in cells lower levels in GDM placentae could imply elevated utilization for placental energy generation. We recently demonstrated that *in-vitro* syncytiotrophoblast rely on FA to support approximately 30% of basal mitochondrial respiration, which increased significantly in male syncytiotrophoblast from GDM placentae which supports our finding of reduced TG which could be an adaptive response to ensure optimal placental function [58].

FA’s are important for normal growth and development of the fetus and their supply is particularly critical in the 3^rd^ trimester of pregnancy [35-37]. Of the total FA’s detected across the 9 lipid classes in placenta, 43 had DHA and 21 had AA, both essential FA’s needed for fetal brain, nervous system development and sourced from maternal blood as they cannot be synthesized in the placenta. Both were reduced in GDM placentae (vs Lean) and associated with TG, PC, PS and PI. Significantly lower levels of placental DHA and AA have been described in obesity and suggested to arise from a defect in biomagnification of LCPUFA [59]. Decreased placental DGLA, but increased AA and DHA in obesity and GDM have also been reported but these studies did not consider fetal sex which may contribute to the discrepancy in findings [60]. We observed a reduction in DHA containing PC, PI and TG in male placentae with GDM whereas female placentae showed a decrease in PC and DG species containing the essential FA’s linoleic, DGLA and AA. These observations broadly agree with findings by O’Neill, K *et al* who in a metabolomic screening of amniotic fluid of in GDM reported a sexual dimorphic effect with decreased medium chain FA’s in a male fetus [61]. Studies on DHA levels in maternal blood and their correlation to cord blood in GDM have reported contrasting results [25] [23] [29]. Discrepancy in type of sample analyzed, plasma vs whole blood, time of sample collection and type of GDM patients (A1 vs A2, diet vs insulin / other medication controlled) considered could all account for these discrepancies. In our well-defined cohort, we did not observe a drastic effect of obesity or GDM or sexual dimorphism in maternal plasma FA composition. Furthermore, there was no overlap in lipid classes or FA’s between maternal plasma and placental samples, nor any drastic differences in maternal plasma. Our observations on reduced placental TG, PC and PI (containing DHA) only in male GDM placentae with no changes in maternal levels could imply their increased breakdown for maintaining optimal transfer to the fetus. We therefore quantified protein expression of MFSD2A - the putative DHA transporter in placenta (Supplementary Fig 2) and did not observe any significant differences. As outlined in Table 1.0, the samples utilized in this study belonged to pregnancies that resulted in appropriate for gestational age (AGA) fetuses. Based on these observations, we propose that the reduced levels of DHA in storage lipids/TG as well as PC observed in male placentae could be a compensatory mechanism to maintain DHA supply to the fetus and maintain its growth trajectory. We acknowledge that low number of samples for male and female fetal samples could be a drawback of the current study, but our overall results warrant for further research into placental lipid composition between AGA and large for gestational age (LGA) fetuses to better understand the effects of maternal obesity and diabetes.

Altogether, this study shows how placental FA distribution changes in a sexually dimorphic manner across different lipid classes with maternal obesity and GDM. As these lipid classes have varied functions in cells and tissue based on their FA composition, the results obtained here imply that obesity and GDM affect placental lipid metabolism in a complicated manner that needs more research.

## DATA AVAILABILITY

The data supporting this study are available at Mass Spectrometry Interactive Virtual Environment (MassIVE; [http://massive.ucsd.edu]) accession number MSV000094962, password: GDM5745. The data will be made public once the manuscript is accepted for publication.

## SUPPORTING INFORMATION

This article contains supporting information.

## ACKNOWLEDGEMENTS

This research is affiliated with the Pacific northwest bioMedical Innovation Co-laboratory (PMedIC) joint research collaboration between OHSU and PNNL, conducted under the Laboratory Directed Research and Development Program at Pacific Northwest National Laboratory, a multiprogram national laboratory operated by Battelle Memorial Institute under Contract No DE-AC05-76RL01830 with the Department of Energy

## AUTHOR CONTRIBUTIONS

LM and KBJ conceptualized the study. EW was involved in sample collection and MV, KS, MM, CN and JK were involved with data generation and analysis. LK and LM were involved with data interpretation and LK wrote the manuscript with contribution from MV, KBJ and LM.

