## Supplementary Figure for "Changes in maternal blood and placental lipidomic profile in obesity and gestational diabetes: Evidence for sexual dimorphism"

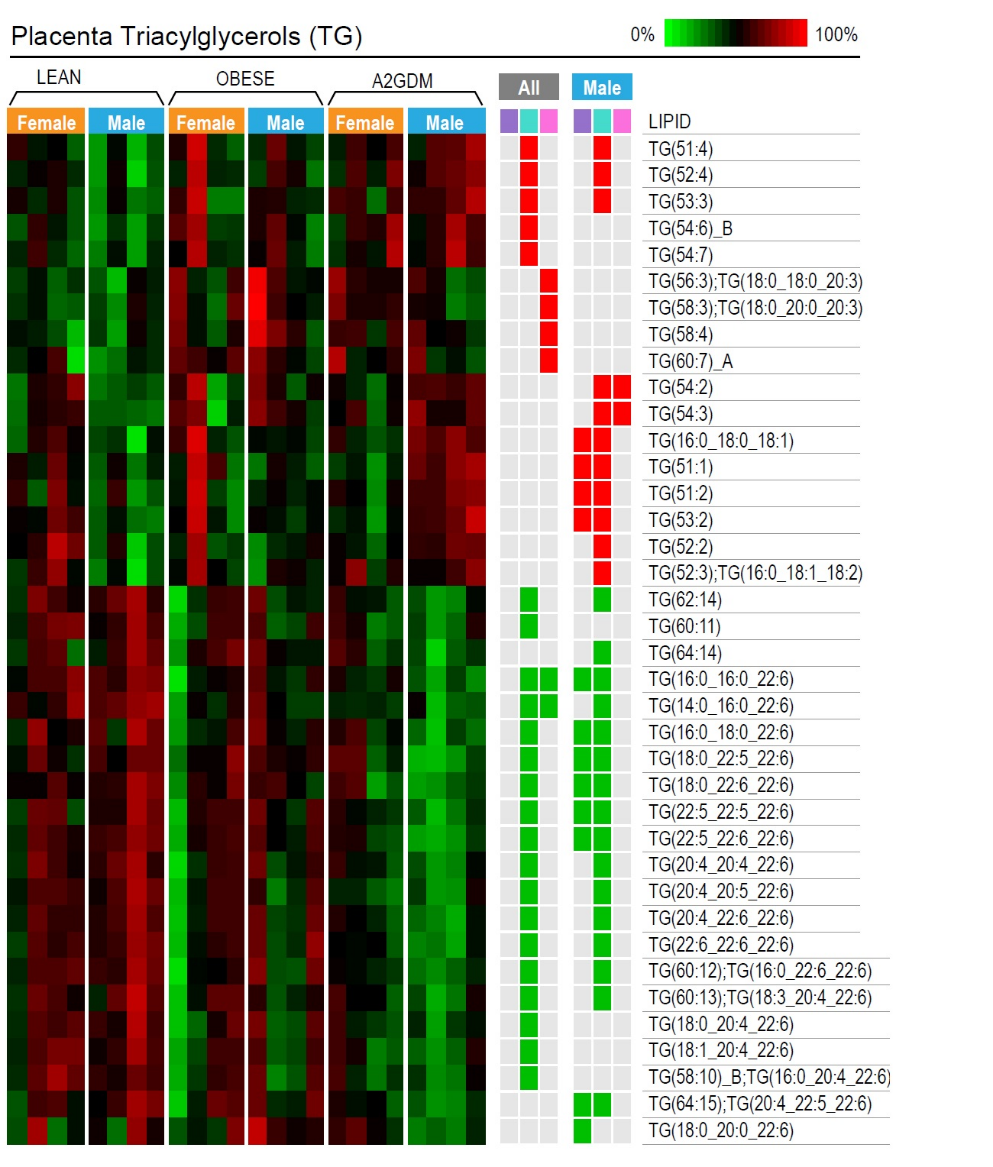

**Supplementary Figure 1: Heatmaps representing the relative log2 expression levels of lipids in Triacylglycerols (TG)** for cases where the TG MS/MS data detected multiple acyl chain identities the TG lipid is label with total number of carbons and double bonds for all three acyl chains. If there were multiple identities, but one was the main species it was listed along with the total number of carbons and double bonds. If there was only one TG species, it is listed alone.

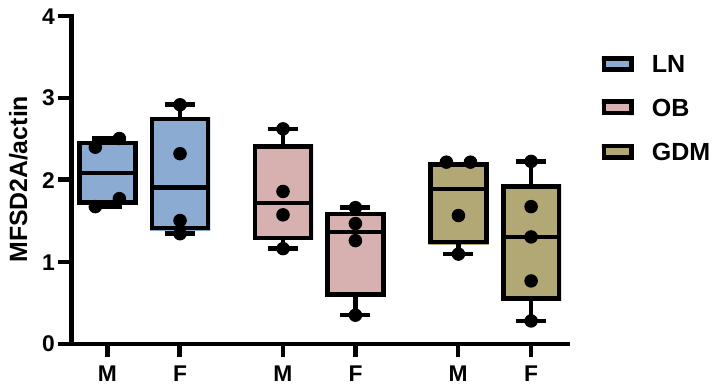

**Supplementary Figure 2:** Protein expression of MFSD2A receptor in male and female placentas from lean, obese and GDM pregnancies (n=5 per group, per sex).
